# Inverse modeling of heterogeneous ECM mechanical properties in nonlinear 3DTFM

**DOI:** 10.1101/2025.02.06.636898

**Authors:** Alejandro Apolinar-Fernández, Jorge Barrasa-Fano, Hans Van Oosterwyck, José A. Sanz-Herrera

## Abstract

Accurate characterization of cellular tractions is crucial for understanding cell-extracellular matrix (ECM) mechanical interactions and their implications in pathology-related situations, yet their direct measurement in experimental setups remains challenging. Traction Force Microscopy (TFM) has emerged as a key methodology to reconstruct traction fields from displacement data obtained via microscopic imaging techniques. While traditional TFM methods assume homogeneous and static ECM properties, the dynamic nature of the ECM through processes such as enzyme–induced collagen degradation or cellmediated collagen deposition i.e. ECM remodeling, requires approaches that account for spatio-temporal evolution of ECM stiffness heterogeneity and other mechanical properties. In this context, we present a novel inverse methodology for 3DTFM, capable of reconstructing spatially heterogeneous distributions of the ECM’s stiffness. Our approach formulates the problem as a PDE-constrained inverse method which searches for both displacement and the stiffness map featured in the selected constitutive law. The elaborated numerical algorithm is integrated then into an iterative Newton-Raphson/Finite Element Method (NR/FEM) framework, bypassing the need for external iterative solvers. We validate our methodology using *in silico* 3DTFM cases based on real cell geometries, modeled within a nonlinear hyperelastic framework suitable for collagen hydrogels. The performance of our approach is evaluated across different noise levels, and compared versus the commonly used iterative L-BFGS algorithm. Besides the novelty of our formulation, we demonstrate the efficacy of our approach both in terms of accuracy and CPU time efficiency.

## 1. Introduction

Mechanobiology is an active research field which investigates the relationship between mechanics and the origin and progression of multiple physiological and pathological processes. Such processes are linked then to cell activity within biological tissues, and fundamental insights on the principles by which mechanical forces and conditions alter the activity of cells and their microenvironment, are of great interest to the fields of biology and medicine [1–3]. Indeed, it has been found that the mechanical interaction between cells and their surrounding extracellular matrix (ECM) guides processess such as bending, stretching, and repositioning of the epithelium, which are required for tissue morphogenesis [4, 5]. These processes also activate signalling pathways that regulate cancer invasion [6], angiogenesis [7, 8], wound healing [9], survival, proliferation and stem-cell differentiation [10–15], and influence other tissue or organ pathologies [16–19]. The mechanism through which cells translate mechanical stimuli into different cellular activities is known as *mechanotransduction*. The intricate nature governing this phenomenom is not yet fully known, and researchers aim to unravel it by means of *in vitro* models that emulate the composition, organization and mechanical characteristics of human-like tissue, which may potentially boost advances in tissue engineering and therapeutic techniques [20–22].

In this context, precise characterization of cellular tractions is essential in the study of mechanotransduction. However, the direct measurement of such tractions and forces in experimental setups is challenging, and several methodologies have been devised and elaborated in the last years [23, 24]. Within these, Traction Force Microscopy (TFM) is emphasized by its effectiveness and potential [25–28]. TFM is a technique that focuses in recovering traction fields associated to specific cell-ECM interactions from measured displacement data obtained via microscopy imaging techniques in the laboratory. The general 3DTFM workflow is summarised as follows: cells are seeded and embedded in a hydrogel that mimics the ECM, which contains spatially distributed fluorescent particles (*beads*). Once cell activity is initiated, a set of microscopy images are captured from which cell morphology and beads motion can be retrieved. In particular, two sets of images are at least needed, one referred to the cell stressed state (reference configuration), and the second corresponding to the cell relaxed state attained after lysis. The force-induced matrix displacement field is then reconstructed by tracking beads motion between the two recorded configurations. Finally, provided the measured displacement field and the constitutive model of the matrix, tractions exerted at the cell-ECM interface are retrieved using the elasticity equations. 3DTFM undoubtedly constitutes a more sophisticated methodology in comparison to its 2D counterpart, providing traction reconstructions in reliable *in vitro* settings that better reproduce the *in vivo* cell environment [29, 30]. 3DTFM presents, however, an increased complexity compared to 2DTFM, regarding experimental setups [6, 8, 31, 32, 32–39], imaging algorithms [26, 33, 40–45], matrix mechanical characterization [27, 39, 40, 46– 49], and computational procedures for traction reconstruction [29, 38, 50–55].

Concerning the computational side of TFM, inverse methods are the most popular methodology for traction reconstruction from measured noisy displacement data. These methods state an optimisation problem in which a new displacement field is searched under the condition that it has to resemble the measured displacements, while being subjected to the fulfilment of fundamental mechanical principles [52]. Inverse methods have been developed over the years and successfully implemented in multiple scenarios. In 2DTFM experiments in which linear elastic behaviour is assumed for the matrix, analytical solutions can be obtained by means of Green’s functions [25]. Regularisation is often applied in order to reduce the impact of noise on the results by penalising high norm values of tractions at certain regions [56]. Regularised inverse approaches have been also employed in 2D/3D linear cases [29, 57–59], linear 3DTFM with constraints [37], and nonlinear 2D/3D with or without constraints [39, 60–62]. In addition, constrained non-regularised methods have been proposed for 3D linear and nonlinear cases [38, 53]. An alternative methodology, the forward method, has also been extensively applied in TFM due to the simplicity of its implementation and its computational efficiency relative to inverse methods [27, 28, 63–65]. In this approach, the measured noisy displacement field is directly interpolated and transformed into a continuous field, which is differentiated to obtain the associated strain field. The latter is introduced in the constitutive equation selected for the matrix to compute the corresponding stresses and tractions. Nonetheless, the accuracy of the forward method highly relies on the quality of the input strains, specially those close to the cell surface, which is often compromised as spatial derivatives tend to amplify the noise contained in the measured displacements. Indeed, it has been shown previously that inverse methods provide substantially more accurate reconstructions than their forward counterpart [38, 50, 51, 53]. An important limitation of the referred works is that these TFM studies assumed the mechanical properties of the ECM as spatially homogeneous and static without considering remodelling. Actually, the ECM is composed of a complex interconnected network of biopolymers, providing structural support for cell activity that serves as a platform for the diffusion of bio-chemicals within tissues. However, the ECM possesses a highly dynamic nature and it undergoes a continuous remodeling process that is crucial for the regulation of diverse cellular behaviors [66]. In particular, cells are able to express enzymes, such as the proteolytic matrix metalloproteinases (MMP), that can locally degrade the collagen microstructure of the ECM [67–72]. MMP-induced degradation facilitates cancer growth and spread by migrating and invading surrounding tissues [73, 74]. In addition, cancer cells deposit siginificant amounts of collagen in their microenvironment, resulting in localized stiffened regions of the matrix [75] that act as walls that shield from drugs that are used during treatments [76, 77]. As a result, the assumption of homogeneity of the ECM properties is not justified in multiple scenarios, therefore compromising the accuracy of traction reconstruction when heterogeneity is disregarded. In fact, the relevance of considering ECM heterogeneity in measuring stresses has been emphasized in several experimental studies [78, 79]. On the other hand, reconstruction of heterogenous ECM patterns may have also a great interest in a number of different biophysical problems [80].

Nonetheless, direct measure of ECM mechanical properties constitutes a difficult challenge, mainly due to cell-induced alterations concentrated in the proximity of cells’ surface [68]. Motivated by this, some authors have proposed innovative inverse formulations to recover material stiffness distributions in 3DTFM experiments. Chen *et al*. [81] implemented a mixed forward/inverse methodology for the reconstruction of ECM stiffness profiles from a measured displacement field. Although interesting, its applicability is limited as it was formulated within the context of linear elasticity. Also, the authors showed stiffness reconstructions directly from measured displacements and, like the majority of forward methods, the associated errors may potentially be non negligible. Song *et al*. [54] proposed a nonlinear inverse formulation for reconstructing ECM stiffness profiles. However, the region in which heterogeneous stiffness values were assumed was small and located in the vicinity of the cell surface. Moreover, the solution approach relies on the use of an external iterative solver, particularly the L-BFGS algorithm, which is potentially inefficient as will be demonstrated in the present study. In addition, they considered a Neo-Hookean model for the ECM, which takes into account geometric nonlinearities due to large deformations, but it does not properly characterize the specific nonlinear behavior of the fibered materials usually employed in 3DTFM. Khan *et al*. [82] implemented a methodology conceptually similar to that of [54], but in this study the region in which heterogeneity can be reconstructed is not limited. The authors modeled the ECM as a Neo-Hookean material as well, and employed the L-BFGS algorithm to solve their formulated inverse problem. Besides, the authors did not verify the accuracy of reconstruction of the proposed methodology with *in silico* cases in which noise in the displacement data is added, as they only assumed noiseless ground truth cases. On the other hand, the study carried out by Mei *et al*. [83], methodologically similar to [54] and [82], outlined a computational methodology for the 2D inverse reconstruction of the shear modulus of cell nuclei (nuclear elastography) by means of a specific version of the adjoint method. They presented an *in silico* TFM study in which they assess on the accuracy of reconstruction of the methodology for different levels of noise added to the ground truth displacements. As in the referred works, the authors employed the iterative solver L-BFGS.

In a previous study, we presented a novel 3DTFM formulation which accounts for MMP-induced ECM degradation [51]. This work establishes a multiscale approach in which 3D degradation profiles are obtained from the solution of a system of partial differential equations. The obtained degradation values are then used to calibrate the parameters of a fiber microstructure generator [84], that fits a continuum (macroscale) nonlinear hyperelastic model in order to consider heterogeneous properties in the 3DTFM workflow. The applicability of this approach is however limited considering that empirical data of ECM degradation are needed to calibrate model parameters of the degradation model. In this framework, we present in this paper a novel methodology to reconstruct spatially heterogeneous distributions of the mechanical parameters of the ECM in a 3DTFM context. The problem is formulated as a PDE-constrained inverse method in which new displacements and the stiffness heterogeneity, are retrieved from the input of the standard measured displacements employed in TFM. The solution to the global constrained problem is approached by iteratively solving two coupled subproblems in which the equilibrium constraint is substituted by a Tikhonov regularization term acting on the hydrogel forces (“weak” enforcement of the equilibrium). Differently to other similar inverse approches, in which iterative methods are used to minimize the cost function, our methodology is developed in the context of a coupled iterative Newton-Raphson/Finite Element Method framework (in which nonlinear elasticity is considered), that does not rely on external solvers. In order to test the methodology, we generated *in silico* 3DTFM cases from a ground truth reference solution, using the geometry of a real cell, and the mechanical behavior of the ECM modeled via a nonlinear hyperelastic law devised for collagen hydrogels. The performance of the proposed methodology is evaluated versus the iterative solver L-BFGS method. Also, we show 3DTFM reconstructions and results corresponding to three different cases referred to the level of noise added to the ground truth displacements solution.

The paper is structured in the following sections: Section 2 presents a detailed description of the mathematical formulation and computational implementation of the proposed inverse methodology. Section 3 illustrates the process of generating the ground truth reference case, the definition of the noise added to the ground truth displacements, and the error metrics used in assessing the accuracy of reconstruction. Section 4 shows and discusses the main results obtained from the study. Finally, the paper ends with some concluding remarks featured in Section 5.

## 2. Mathematical formulation

In this section, the mathematical and computational aspects of the method are presented in detail. The aim of the formulation is to reconstruct the heterogeneous stiffness profile *κ*_*o*_(*x, y, z*) by means of a PDE-constrained inverse method, in the context of a FE framework in which the equilibrium condition is imposed. The parameter *κ*_*o*_ is the one associated to the stiffness of the material in the nonlinear hyperelastic law selected to reproduce the ECM’s mechanical behavior (Section 3.2). The main inverse problem is divided into two coupled subproblems that are solved iteratively similarly to a fixed point procedure. Both subproblems contain a Tikhonov regularization term that serves as a *weak* form of imposing the equilibrium condition. The two subproblems are integrated and solved within a Newton-Raphson/Finite Element Method (NR/FEM) formulation.

The global problem is written to find both the new displacement field **u** and the heterogeneous stiffness distribution *κ*_*o*_ so that the former resemble the measured displacement field **u**^∗^ and the latter behaves smoothly, provided that the solution to the problem satisfies the equilibrium of forces,

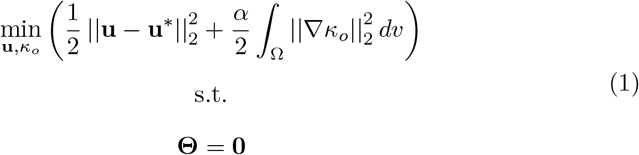

where *α* is the regularization parameter, Ω represents the hydrogel domain and Θ the equilibrium constraint manifold in which the candidate solution must lie.

Normally, in order to solve this problem, the equilibrium contraint would be introduced into the cost function through a Lagrange multiplier field *η* in the following way,

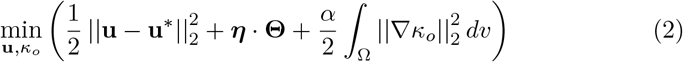

The methodology presented in this paper is based on the substitution of the “strong” constraint, represented by ***η*** *·* **Θ**, by a term that regularizes and minimizes the nodal forces within the hydrogel domain. We refer to this as a “weak” imposition of the equilibrium condition, and it is motivated to keep the numerical workflow of the algorithm as will be seen in the next sections.

The global problem (2) is subdivided into two additional coupled problems that are solved simultaneously: the *forward* and *inverse* subproblems, associated to the terms 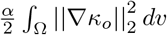 and 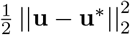, respectively; both approximating the condition represented by ***η*** *·* **Θ** with the aforementioned regularization term. The details about the implementation of both subproblems are described thoroughly in the next sections.

### 2.1 Forward subproblem

The *forward subproblem* is formulated for a given displacement field **u** as input data. As in a conventional forward method, this displacement field is fixed during the solution of this subproblem. At the first global iteration, which involves the forward and inverse subproblems, its value is fixed to the measured displacements field **u**^∗^. The problem (2) is then stated as,

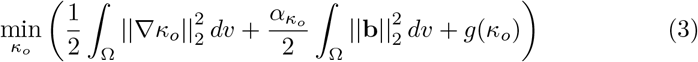

in which 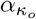 is a regularization parameter associated to the forward subproblem, and **b** represents the body forces acting in the ECM’s interior domain. Overall, this regularization term establishes the equilibrium condition through 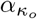 as **b** must vanish within the hydrogel domain. *g*(*κ*_*o*_) is a sigmoid (activation) function included to avoid negative values of *κ*_*o*_ during the iterative process.

The problem stated in Eq. (3) has a minimum stationary solution at,

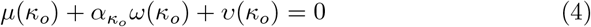

where,

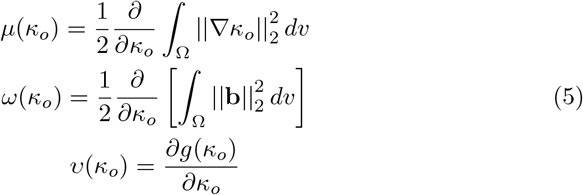

Eqs. (5) are developed into their discretized form within a Finite Element (FE) framework. In this sense, field variables are discretized as *κ*_*o*_ ≈ **N**^*e*^ *·* ***κ***_***o***_^*i*^, ∆*κ*_*o*_ ≈ **N**^*e*^ *·* ∆***κ***_***o***_^*i*^, **b** ≈ **N**^*e*^ *·* **b**^*i*^ and *g* ≈ **N**^*e*^ *·* ***g***^*i*^, where **N**^*e*^ is the matrix that contains the shape functions of the finite element (*e*). Further, ***κ***_***o***_^*i*^, ∆***κ***_***o***_^*i*^, **b**^*i*^ and ***g***^*i*^ are discrete vectors which contain the discrete values of *κ*_*o*_, ∆*κ*_*o*_, **b** and *g* at node positions *i*, respectively, in the FE element (*e*). Introducing this discretization into Eqs. (5) yields,

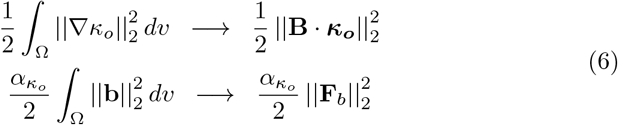

**B** is the global gradient matrix after assembly, and ***κ***_***o***_ the global vector with discrete values of *κ*_*o*_ at each node of the FE mesh. **F**_*b*_ is the global vector, after assembly, with discrete values corresponding to the body forces in the internal ECM domain.

Functions *μ, ω* and *υ* in Eq. (5) now take the discretized expressions,

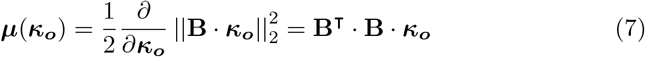

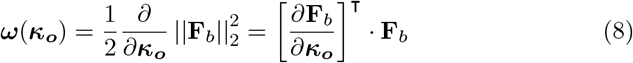

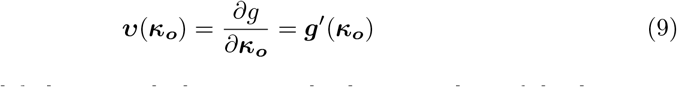

with ***g***^*′*^ a nodal global vector which contains the discrete values of the derivative of function *g* evaluated at ***κ***_***o***_. These equations are linearized as follows:

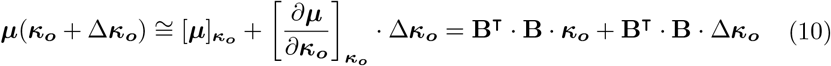

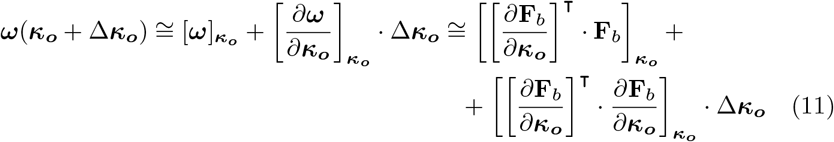

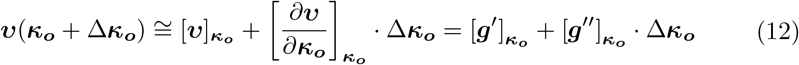

with ***g***^*′′*^ a nodal global vector which contains the discrete values of the second derivative of function *g* evaluated at ***κ***_***o***_. Note that the second derivative of **F**_*b*_ with respect to ***κ***_***o***_ was neglected in Eq. (11). On the other hand, using the spatial version of the discretized principle of virtual work (PVW), the following relation can be written

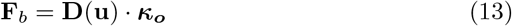

Global matrix **D** is obtained after assembly of the PVW which depends on the displacements, which are known and fixed in this forward subproblem. Note that **F**_*b*_ is linearly related with ***κ***_***o***_ since stresses are linear with *κ*_*o*_ as is seen in the selected hyperelastic law (see Section 3.2).

Using Eq. (13) in Eqs. (10), (11) and (12); Eq (4) yields,

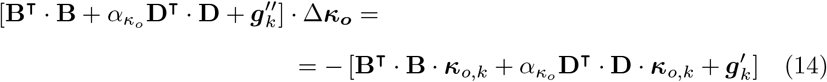

Remarkably, nonlinearity is introduced in Eq. (14) through the activation function *g*. Therefore, this equation is iteratively solved until convergence is achieved. Note that vector ***κ***_*o*_ is defined in the nodes of the mesh according to the developed FE formulation. As a result, it is mathematically impossible to fulfill the strong form of the equilibrium condition as it would involves the fulfillment of 3 equations per node with just one unknown variable (the stiffness) per node. Consequently, this condition is imposed in our formulation through parameter 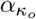 in Eq. (14).

### 2.2 Inverse subproblem

The *inverse subproblem* is formulated under the assumption that *κ*_*o*_ is known and fixed, as it results from the forward subproblem. Hence, the problem is stated as a normal unconstrained inverse problem with regularization in which displacements are recomputed and the material properties are known (although inhomogeneous). The problem can be written as follows,

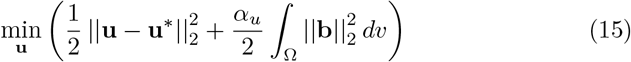

in which *α*_*u*_ is the regularization term parameter associated to the inverse subproblem, and **b** is defined analogously to the forward subproblem.

Similarly to the previous section, Eq. (15) has a minimum stationary solution using the Gateaux derivative with respect to **u**,

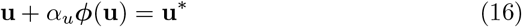

where ***ϕ***(**u**) is defined as a nonlinear function that depends implicitly on **u** through **b**,

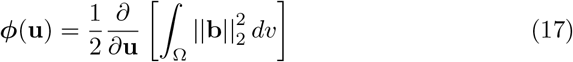

***ϕ*** is linearized in the vicinity of **u** in the following fashion,

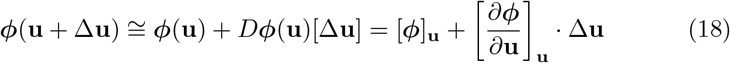

being *D****ϕ***(**u**)[∆**u**] the directional derivative of *ϕ*(**u**) in the direction of an arbitrary small increment ∆**u**. The linearized version of Eq. (16) is then written as,

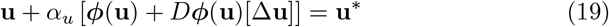

which will be discretized following a Finite Element approach and solved iteratively using a Newton-Raphson procedure. The reader is addressed to Appendix A of a previous paper [52] for further details of a similar problem.

As in the forward subproblem, the field variables are discretized such that **u** ≈ **N**^*e*^ *·* **u**^*i*^, ∆**u** ≈ **N**^*e*^ *·* ∆**u**^*i*^ and **u**^∗^ ≈ **N**^*e*^ *·* **u**^∗,*i*^. Besides, **u**^*i*^, ∆**u**^*i*^ and **u**^∗,*i*^ are vectors whose components correspond to the discrete values of **u**, ∆**u** and **u**^∗^ at node positions *i*, respectively, in the FE element (*e*). Finally, after assembly, the resulting linearized global system yields the expression,

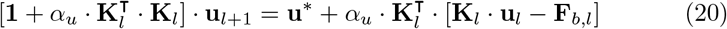

where **1** is the unit matrix, **u**_*l*+1_ and **u**_*l*_ are the vectors corresponding to the nodal values of the reconstructed displacements at the current *l*+1, and previous *l* iterations, respectively. **F**_*b*,*l*_ and **K**_*l*_ are the global vector of internal forces and the tangent stiffness matrix, respectively, both evaluated at the *l*-th iteration. The coupled algorithm of solving both the forward and inverse subproblems within the global iterative procedure is explained in the next section.

### 2.3 Global iteration description

Both forward and inverse subproblems are solved sequentially within a global iterative loop that is stopped when the global convergence criterion 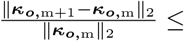 TOL is achieved (*m* being the counter of the global loop). On the one hand, the forward subproblem iterates with respect to a particular local convergence criterion defined for 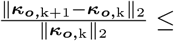 TOL (*k* being the local counter of iterations of the forward subproblem). On the other hand, the inverse subproblem proceeds up to convergence of 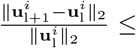 TOL ( being the local counter of iterations of the inverse subproblem). TOL is the relative convergence tolerance, which is set as 1e-5 in this work. A schematic representation of this process is illustrated in Figure 1.

**Figure 1:**
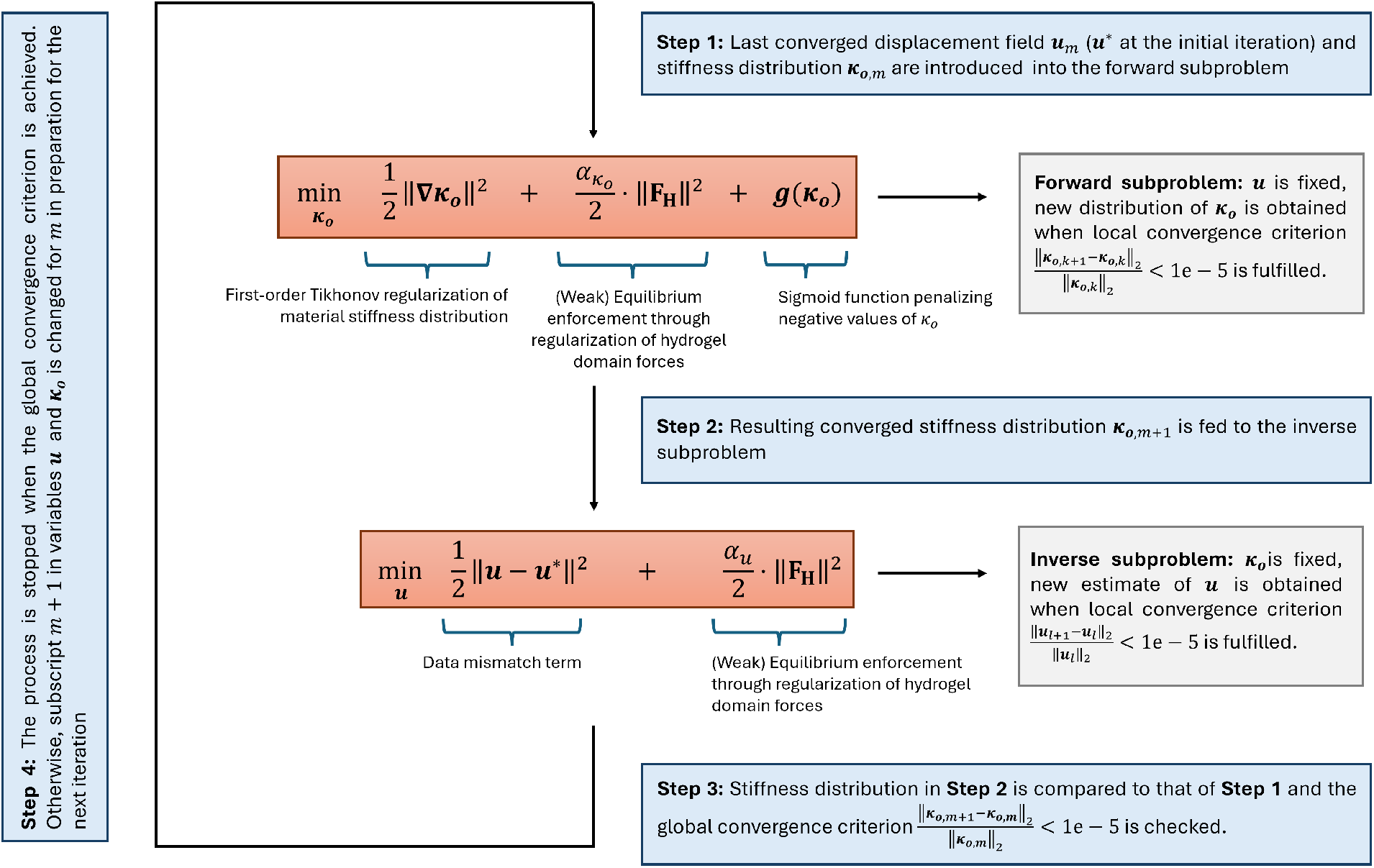
Workflow of the iterative process: Convergence of the forward subproblem, provided a fixed displacement vector **u**, yields the discrete stiffness vector ***κ***_***o***_, which is the input quantity of the inverse subproblem. Convergence of the inverse subproblem, provided a fixed ***κ***_***o***_, yields the discrete displacement vector **u**, which is the input quantity of the forward subproblem at the next global iteration. The global iterative procedure is stopped when the global convergence is achieved.

### 2.4 Calibration of the regularization parameters

The “weak” enforcement of the equilibrium condition through regularization in the forward and inverse subproblems yields regularization parameters 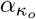 and *α*_*u*_. The calibration procedure for both parameters is based on a modified version of the frequently used L-curve method [60, 85, 86]. First, in the forward subproblem, 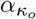 is taken as large as possible to enforce equilibrium while at the same time keeping the reconstructed stiffness values in the non-negative range. Once 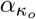 is calibrated under this criterion, a sweep of simulations for different values of *α*_*u*_ is performed to reproduce the L-curve. In order to do so, the regularization term 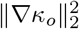 is represented against the data mismatch term 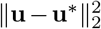 for the simulated values of *α*_*u*_. Then, the optimal value is determined as the one that is closest to the point of maximum curvature of the resulting curve. The reader is addressed to [52] for a more detailed description of the L-curve method applied to the inverse subproblem. Note that *α*_*u*_ is the only one parameter calibrated with the L-curve method, and it is not required to be calibrated for multiple values of 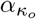 .

## 3. Setup of the synthetic 3D TFM cases

This section describes the process for generating the ground truth case that will be taken as a reference for the assessment of the validity of the results. The displacements required as an input in the methodology are then taken from this ground truth case. First, information about the real 3DTFM experiment, the employed cell geometry, as well as details about the nonlinear hyperelastic model selected to represent the mechanical behavior of the hydrogel are given. Afterwards, the configuration of the ground truth scenario, and the process of generating the input displacement fields for the *in silico* TFM reconstructions are explained. Finally, the error metrics used for the assessment of the performance of the methodology are introduced.

### 3.1 Acquisition of microscopy images, segmentation and generation of the FE mesh

The *in vitro* setting from which the geometry of the cell considered for this study was extracted is outlined in previous papers published by our lab [52, 87]. In this section, a brief description of the generation process of the finite element mesh corresponding to the cell geometry concerned is given.

The selected cell geometry corresponds to a human breast cancer cell (line MDA-MB-231). For the associated experiment, the cell was embedded in a 1.2 mg/ml collagen-based hydrogel prepared with a mix of rat tail and bovine skin collagen, following the protocol described in [31]. 24 hours after polymerization, confocal images of the cell were captured, featuring a voxel size of 0.227 x 0.227 x 1.48 μm^3^. The obtained images were then enhanced by a contrast stretching operation and subsequently segmented, resulting in a 3D binarized image of the cell geometry. Finally, the MatLab toolbox Iso2Mesh [88] was employed for the meshing of the voxelized cell boundary geometry resulting from the segmentation process. The finite element mesh is composed of 4-noded tetrahedra with linear interpolation, resulting in a total of 34666 nodes and 199691 elements, which includes the hydrogel domain and the cell in its center (Figure 2a).

**Figure 2:**
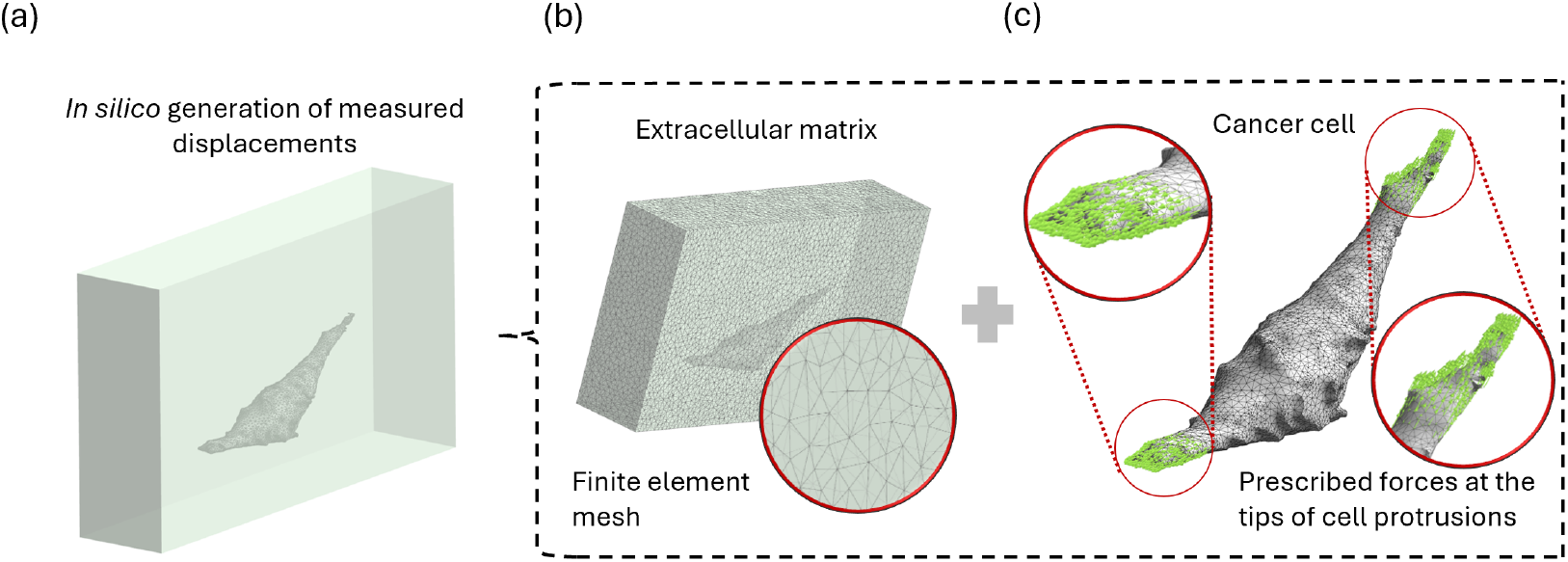
Setup of the ground truth reference case: (a) Problem domain containing both the hydrogel and the cell. (b) 3D tetrahedral Finite Element mesh. (c) Ground truth forces (arrows) prescribed on the tips of the cell’s protrusions.

### 3.2. Mechanical characterization of the hydrogel

The mechanical characterization of the ECM-mimicking collagen hydrogel is carried out by fitting the model parameters to a specific nonlinear hyperelastic model in shear rheology experiments. The selected nonlinear model was used in previous publications by the authors [50–52], and it was originally presented by Steinwachs *et al*. in [39], where it was proven to be an excellent model to reproduce the mechanical behavior of collagen hydrogels. Specifically, the model considers the mechanical behavior of a single fiber, that can be subjected to three different states of deformation: buckling (*λ <* 0), in which the fiber is unstable and its stiffness decay exponentially; straightening (0 ≤ *λ < λ*_*s*_), in which the fiber recovers its natural length (*λ*_*s*_) as it straightens, being the stiffness kept as constant; and stretching (*λ* ≥ *λ*_*s*_), in which the fiber stretches further than its natural length and its stiffness increases exponentially. *λ* denotes the macroscopic stretch of a fiber. Accordingly, the strain energy density function associated to a single fiber is defined from the stiffness through its second derivative with respect to *λ*,

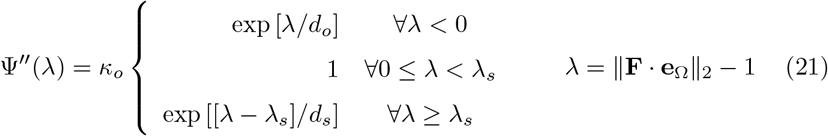

where *κ*_*o*_ is the material stiffness, *d*_*o*_ is the non-dimensional buckling parameter, *d*_*s*_ the non-dimensional strain-stiffening (dimensionless) parameter, and *λ*_*s*_ the non-dimensional parameter that defines the extension of the interval within which the fiber behaves linearly, i.e. with constant stiffness. Moreover, **e**_Ω_ is the unit vector that represents the fiber’s orientation and **F** is the deformation gradient tensor. The macroscopic behavior of the hydrogel is obtained by averaging fiber contributions to the strain energy in all directions. Thus, the constitutive law in mixed configuration is obtained as,

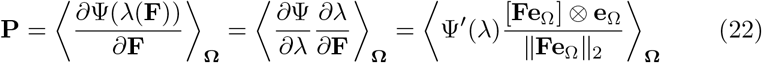

where **P** denotes the First Piola-Kirchhoff stress tensor and the brackets indicate the average over all directions Ω in the unit sphere.

The model is integrated into the NR/FEM computational framework described in Section 2 by means of an ABAQUS UMAT subroutine. The reader is referred to the Appendices in [50] where a thorough description of the model formulation and its computational implementation is presented.

For this study, the parameters in Eq. (21) were fitted to reproduce the shear rheology curves obtained from real collagen hydrogels in the laboratory shown in a previous paper by the authors [52]. The values of the parameters resulting from the fitting process are given in Table 1. These values correspond to the virgin state (non-degraded, homogeneous) of the ECM.

**Table 1:**
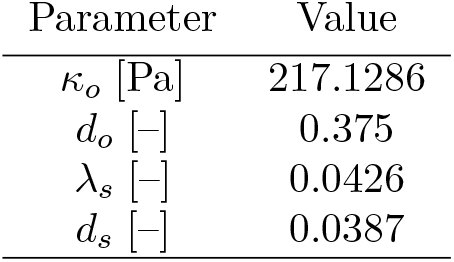
Virgin (non-degraded, homogeneous ECM) values of the nonlinear model parameters (Eq. 21).

| Parameter | Value |
| --- | --- |
| $\kappa_o$ [Pa] | 217.1286 |
| $d_o$ [-] | 0.375 |
| $\lambda_s$ [-] | 0.0426 |
| $d_s$ [-] | 0.0387 |

#### 3.3 Generation of the ground-truth cases

The aim of the methodology presented in this paper is to reconstruct stiffness profiles from standard input displacement data in the context of TFM. Therefore, different heterogeneous stiffness profiles *κ*_*o*_ are prescribed in the hydrogel domain of the ground truth reference case presented in Figure 2a. The FE mesh of this case of study is shown in Figure 2b. These profiles simulate degradation effects of the substances secreted by the cell and remodelling on the collagen hydrogel microstructure. In particular, two different stiffness heterogeneities were simulated, hereby referred to as tip-localized matrix degradation pattern (first pattern) and diffuse matrix remodeling pattern (second pattern).

The tip-localized degradation pattern is generated as an exponential decay, following a sigmoid-type function in the hydrogel domain from the emission spot that is located at the tip of one of the cell’s protrusions. Figure 3a shows a view of the mid cross-section of the hydrogel domain in which the referred ground truth degradation pattern is outlined. Figure 3b illustrates a projection of the cross-section on the XY-plane, and Figure 3c plots the ground-truth stiffness profile with respect to the distance from the emission point (*γ*) along the green line represented in Figure 3b. Secondly, the diffuse matrix remodeling pattern is shown in Figure 4. Differently to the tip-localized matrix degradation pattern, this one outlines the process of matrix remodeling which induces stiffening of the ECM around the vicinity of the cell’s surface. This can be attributed to cellmediated collagen deposition process, followed by matrix degradation. Then, nominal (virgin) hydrogel stiffness is assumed far from the cell location. This kind of stiffness distribution is typically seen in *in vitro* setups [75–77].

**Figure 3:**
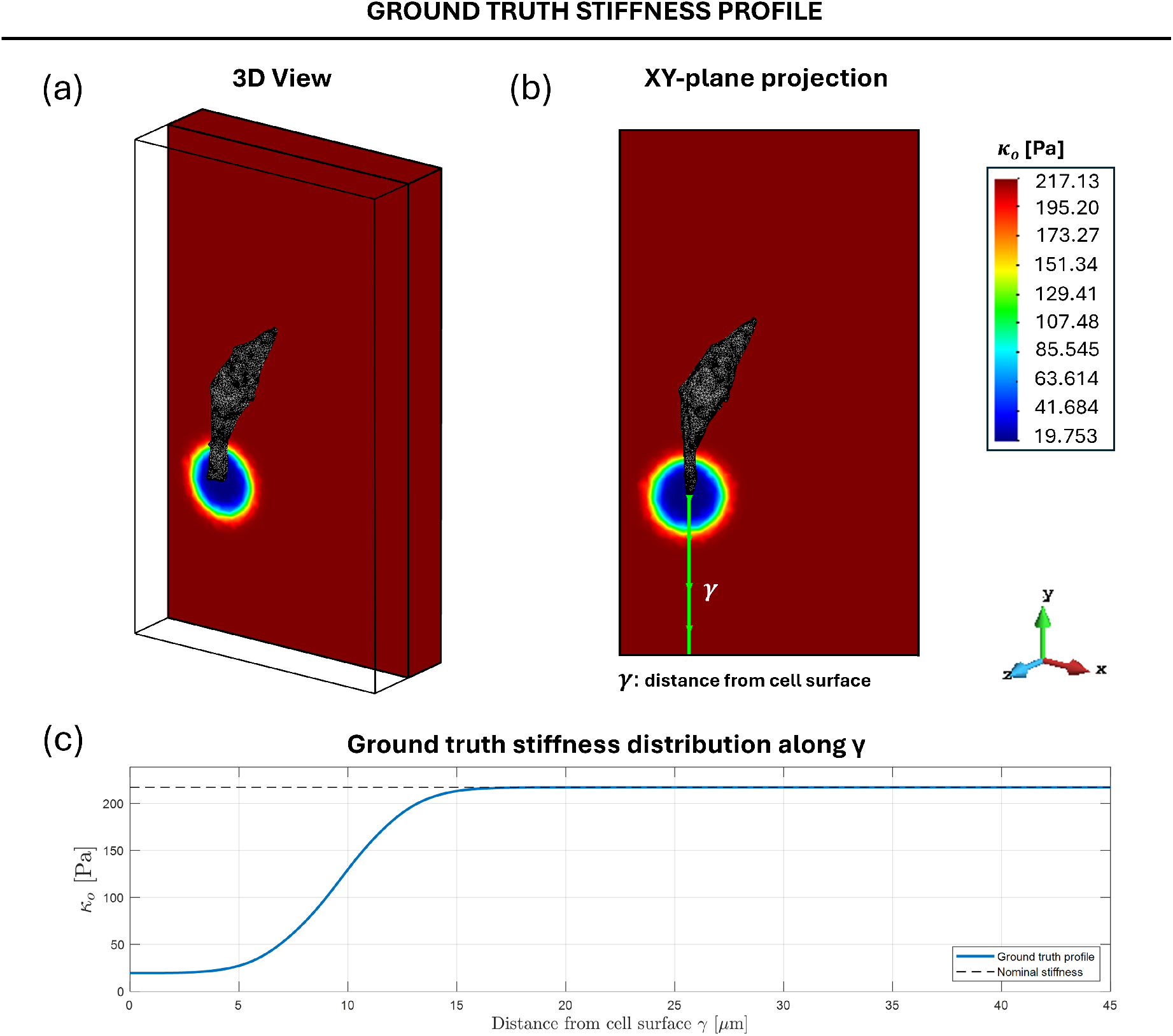
Material stiffness *κ*_*o*_ contours for the ground truth case (tip-localized matrix degradation pattern): (a) 3D view of mid cross-section of the hydrogel domain outlining spatial material stiffness distribution. (b) XY-plane projection of the mid cross section. (c) Stiffness plotted with respect to the distance from the emission point (cell’s tip) highlighted as a green line drawn in subfigure (b).

**Figure 4:**
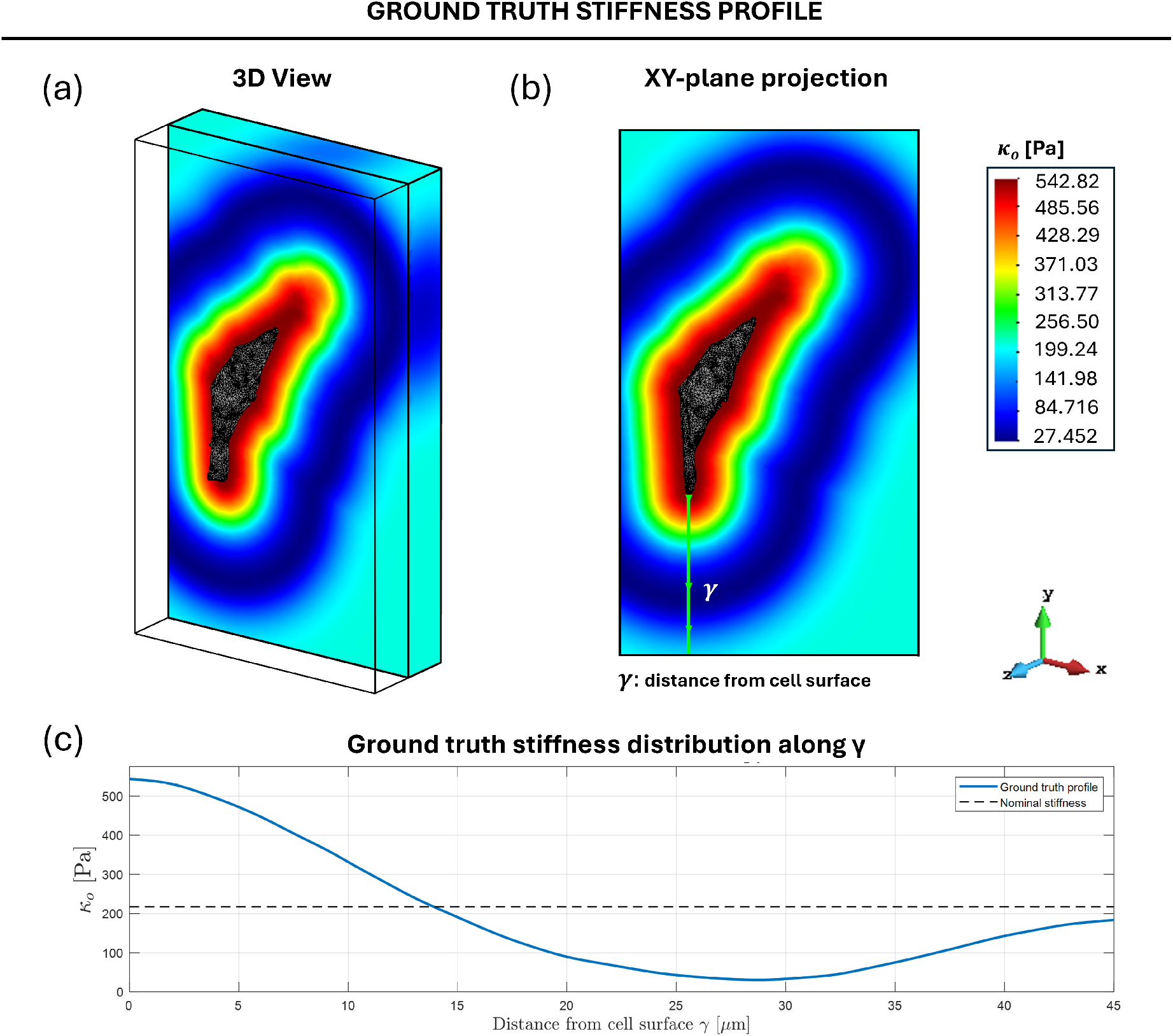
Material stiffness *κ*_*o*_ contours for the ground truth case (diffuse matrix remodeling pattern): (a) 3D view of mid cross-section of the hydrogel domain outlining spatial material stiffness distribution. (b) XY-plane projection of the mid cross section. (c) Stiffness plotted with respect to the distance from the emission point (cell’s tip) highlighted as a green line drawn in subfigure (b).

Afterwards, for the two considered stiffness profiles, nodal forces are prescribed on the tips of the cell’s protrusions and null normal displacement boundary conditions are applied to the hydrogel boundaries. The values of the forces were calibrated such that the magnitude of the associated deformations is aligned with our observations in the laboratory. Figure 2c illustrates the regions where these cellular forces are prescribed. The field variables of interest (stresses, strains, tractions, displacements) are retrieved from the solution to the corresponding nonlinear elasticity problem simulated by means of the FE software Simulia ABAQUS, and using the described mechanical behavior of the hydrogel through an external UMAT subroutine. Together with the material stiffness distribution, the recovery of these variables using the presented inverse methodology, are compared versus the ground truth solution for the different levels of noise considered in the study.

The recovery of the ground truth solutions corresponding to the 2 stiffness patterns (Figures 3 and 4) are compared using our proposed methodology versus the frequently used L-BFGS iterative algorithm (Section 4.1).

### 3.4. Input displacement fields and error metric

In order to generate the input data of the inverse algorithm described in Section 2, the ground truth displacements (obtained as explained in Section 3.3) are “corrupted” by adding a Gaussian noise to the value of each degree of freedom of the FEM-discretized ground truth displacement vector,

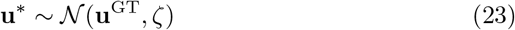

with **u**^∗^ being the measured (simulated) displacement vector and **u**^GT^ the ground truth displacement vector. *𝒩* (**u**^GT^, *ζ*) is a normal distribution with mean the values of vector **u**^GT^, and *ζ* its standard deviation with respect to the ground truth displacements. Three different levels of noise have been considered for assessment: 2% (*ζ* = 0.000292), 5% error (*ζ* = 0.000741) and 10% (*ζ* = 0.001475) error in displacements .

On the other hand, in order to quantify the accuracy of the method in regards to material stiffness reconstruction at the different noise levels employed in this study, a specific error metric is defined as follows:

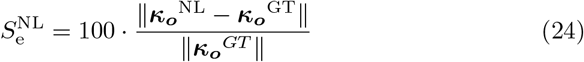

where ***κ***_***o***_^NL^ represents the vector of the (interpolated) stiffness at hydrogel points along the reference line (*γ*) shown in Figure 3b for a certain noise level (NL), and ***κ***_***o***_^*GT*^ being the corresponding ground truth case.

## 4. Results and discussion

In this section, the main results of the study are presented. First, the proposed methodology is compared with an iterative algorithm usually employed in the context of the application on hand. After this, the results corresponding to the recovery of variables performed for synthetic 3DTFM cases are presented, for three different levels of noise added to the ground truth solution. The resulting data were post-processed in different ways in order to assess the accuracy that the methodology provides. In particular, its performance is evaluated based on the accuracy of reconstruction for stiffness, tractions and displacements. In terms of computer performance, the number of iterations taken in each case, until local and global convergence are achieved, are discussed. Both local convergence and the global convergence criteria were established as a relative change (tolerance) between iterations of 1e-5 (as described in Section 2.3).

### 4.1. Noise-free comparison between NR/FEM and L-BFGS algorithms

In order to establish a comparison between the proposed methodology and the frequently used L-BFGS iterative algorithm [54, 82, 83], noise-free recovery of ground truth solutions are performed. L-BFGS algorithm can be quite straightforwardly implemented as it just requires the cost function (Eq. (3)) and its gradient (Eq. (4)).

Two degradation patterns are considered for reconstruction given in Figures 3 and 4. As displacements are considered as noise-free, just the forward subproblem is required for the comparison between solvers. Figures 5 and 6 show the reconstructed ground truth stiffness profiles for the tip-localized matrix degradation and diffuse matrix remodeling patterns, respectively, using the L-BFGS iterative solver (Figures 5a and 6a), and the NR/FEM proposed methodology (Figures 5b and 6b). These results are qualitatively compared to the ground truth solution in Figures 5c and 6c; and quantitatively in (Figures 5d and 6d). On the other hand Table 2 provides, for each methodology, the values of the defined performance metrics for reconstruction of each degradation pattern. Qualitatively, it can be observed in Figures 5 and 6 that both methodologies are able to successfully recover the ground truth solution, being the NR/FEM reconstruction overall better than the L-BFGS solver. Nevertheless, Table 2 reveals that, although L-BFGS takes a tenth of the time per iteration that NR/FEM, as no inversion of any matrix is required in this algorithm, the number of total iterations is significantly bigger for the L-BFGS. In fact, the predefined number of iterations (2000) was achieved before the predefined tolerance for convergence (the same for both methodologies) was reached for the tip-localized pattern. As a result, the total time of execution of L-BFGS relative to NR/FEM is nearly sixfold in the case of the tip-localized matrix degradation pattern, and around threefold in the case of the diffuse matrix remodeling pattern. Also, it has to be taken into account that these times of execution correspond to the forward subproblem. The full iteration procedure (both forward and inverse subproblems, see Figure 1) is required when noisy displacement fields are considered. Therefore, the tangent stiffness matrix has to be updated and assembled with the new displacement field during iterations. As a consequence, given the discrepancy between the number of iterations performed by L-BFGS and NR/FEM, the total CPU time will heavily increase, highlighting that the proposed NR/FEM scheme outperforms versus the L-BFGS iterative solver.

**Table 2:**
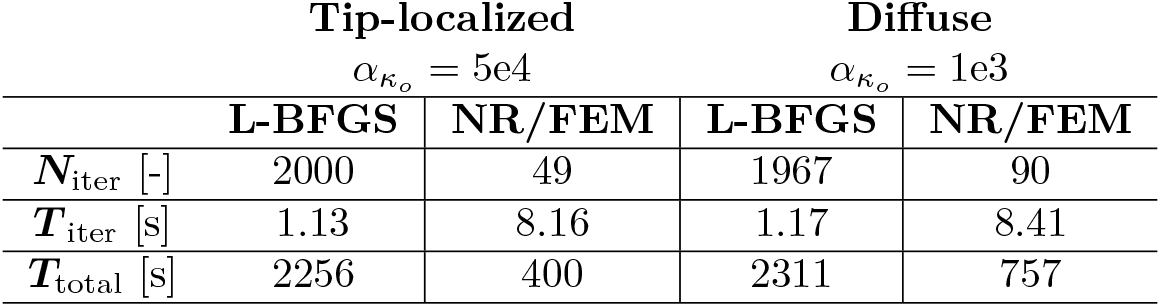
Performance metrics for both the L-BFGS algorithm and the proposed methodology introduced in this paper (NR/FEM) (in downward direction): Number of iterations, CPU time per iteration and total CPU time taken for the reconstruction. The calibrated regularization parameters of the forward subproblem 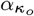 are given for each degradation pattern.

**Figure 5:**
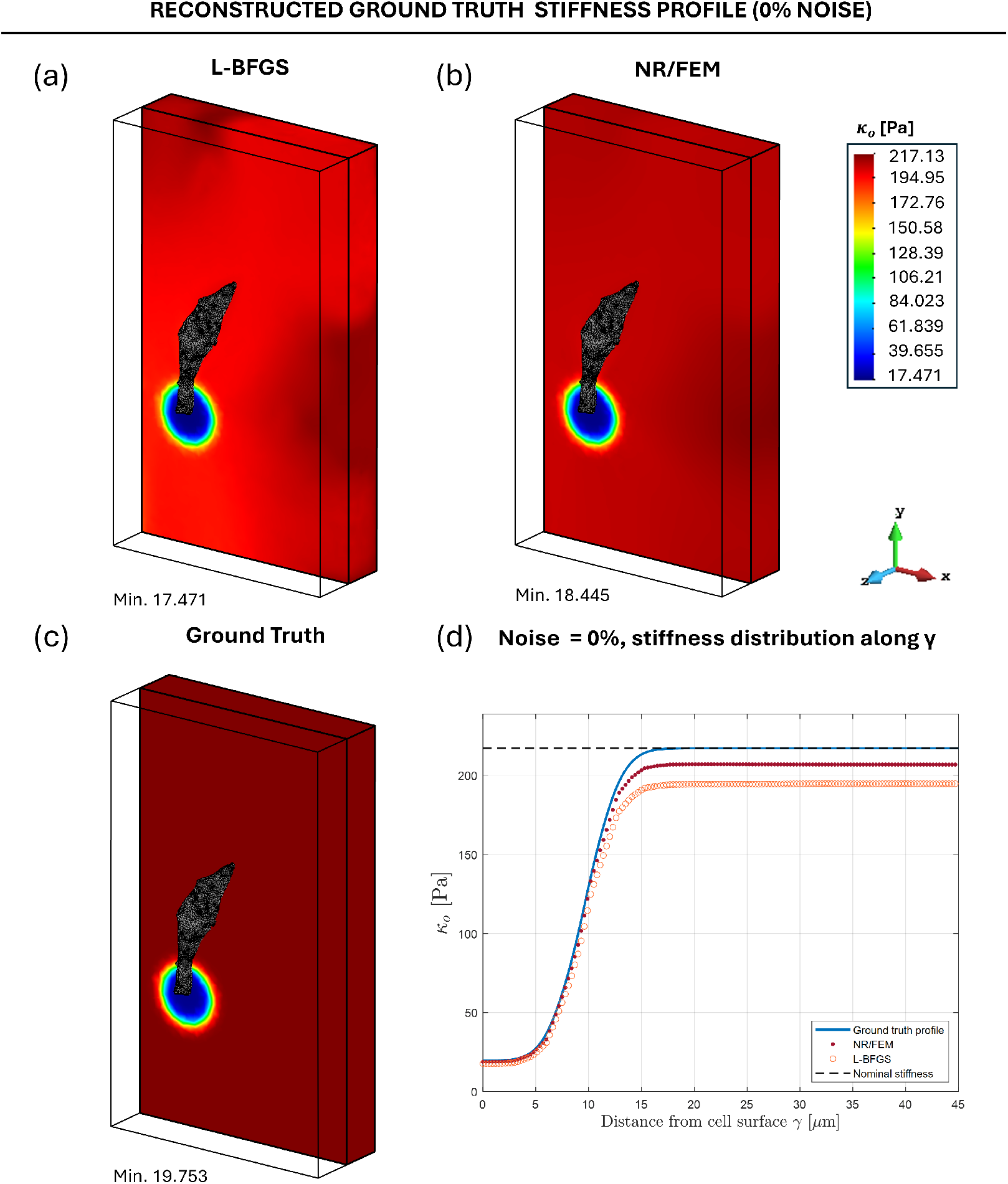
Material stiffness *κ*_*o*_ contours for the reconstructed ground truth case (tip-localized matrix degradation pattern): (a) 3D view of mid cross-section of the hydrogel domain outlining reconstructed spatial material stiffness distribution (L-BFGS). (b) 3D view of mid cross-section of the hydrogel domain outlining reconstructed spatial material stiffness distribution (NR/FEM). (c) Ground Truth spatial material stiffness distribution. (d) Comparison of reconstructed stiffness profiles with the ground truth profile along *γ* (see Figure 3).

**Figure 6:**
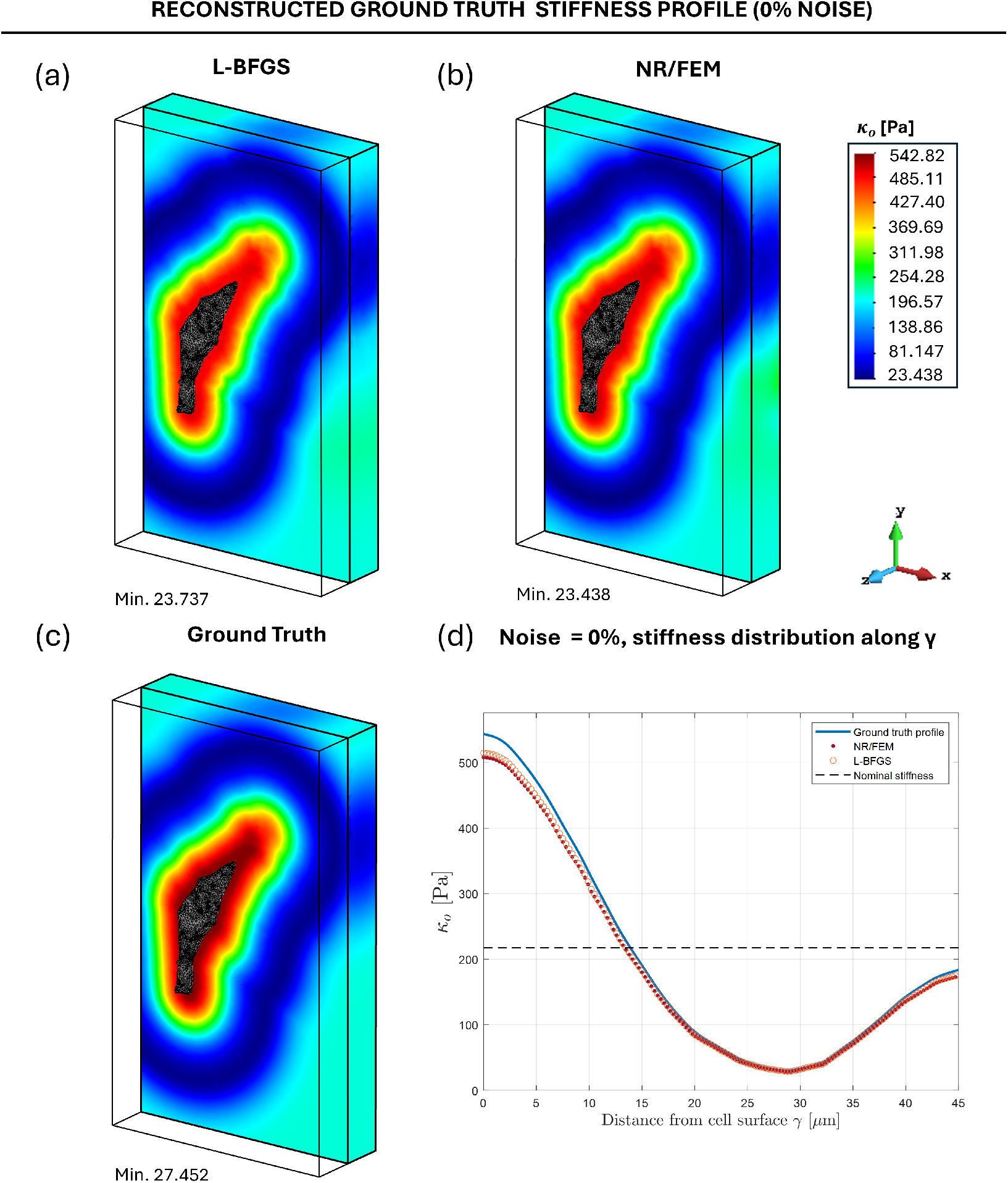
Material stiffness *κ*_*o*_ contours for the reconstructed ground truth case (diffuse matrix remodeling pattern): (a) 3D view of mid cross-section of the hydrogel domain outlining reconstructed spatial material stiffness distribution (L-BFGS). (b) 3D view of mid cross-section of the hydrogel domain outlining reconstructed spatial material stiffness distribution (NR/FEM). (c) Ground Truth spatial material stiffness distribution. (d) Comparison of reconstructed stiffness profiles with the ground truth profile along *γ* (see Figure 4).

### 4.2. Results of reconstruction with noisy displacements

Results of reconstructed variables, following the proposed methodology introduced in Section 2, are given in this section. Different levels of noise in the input displacements are considered, using Eq. (23), to the tip-localized degradation pattern case. Figures 7, 8 and 9 show a visual representation of the global reconstructed stiffness profiles for noise levels 2%, 5% and 10%, respectively. The ground truth solution is also included in these figures for comparison purposes. It can be observed in Figures 7a, 8a and 9a that, in general terms, the recovered profiles are similar to that of the ground truth case. Moreover, it is shown in Figures 7c, 8c and 9c that the stiffness distribution progressively struggle to capture the values located in the vicinity of the emission point on the cell protrusion as noise level increases. Moreover, Table 3 quantitatively provides the error metric *S*_*e*_ (Eq. (24)) of the stiffness reconstruction, showing that this error steadily increases from 5.31 to 8.56%, when the added noise in the displacement fields increases from 2% to 10%. In addition, Table 3 provides the number of global and local iterations required for achieving convergence of the method. It can be seen that, as noise level increases, the convergence of the iterative processes becomes more difficult, requiring generally more iterations. Particularly, the amount of both local iterations and that of the global iterations steadily increase with noise level, this tendency being substantially emphasized for the 10% noise level case. In the latter, the number of iterations required for the convergence of the inverse subproblem rises significantly compared to the increase observed in the other cases.

**Table 3:** Values of the defined error metrics (stiffness, percentages) for each noise level case added to the displacement field of the tip-localized degradation pattern case. The last three columns contain, respectively, the values of the number of global iterations, local iterations of the forward subproblem and local iterations of the inverse subproblem. The values of 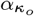and *αu* correspond to the optimal regularization parameters obtained via the modified L-curve method described in Section 2.4.

| | $\alpha_{\kappa_o}$ | $\alpha_u$ | $S_e$ | $N_{\text{iter},m}$ | $N_{\text{iter},k}$ | $N_{\text{iter},l}$ |
| --- | --- | --- | --- | --- | --- | --- |
| **2% noise** | 1e2 | 5e-3 | 5.31 | 13 | 65 | 24 |
| **5% noise** | 1e2 | 1e-2 | 6.68 | 20 | 98 | 39 |
| **10% noise** | 1e2 | 1e-2 | 8.56 | 37 | 168 | 85 |

**Figure 7:**
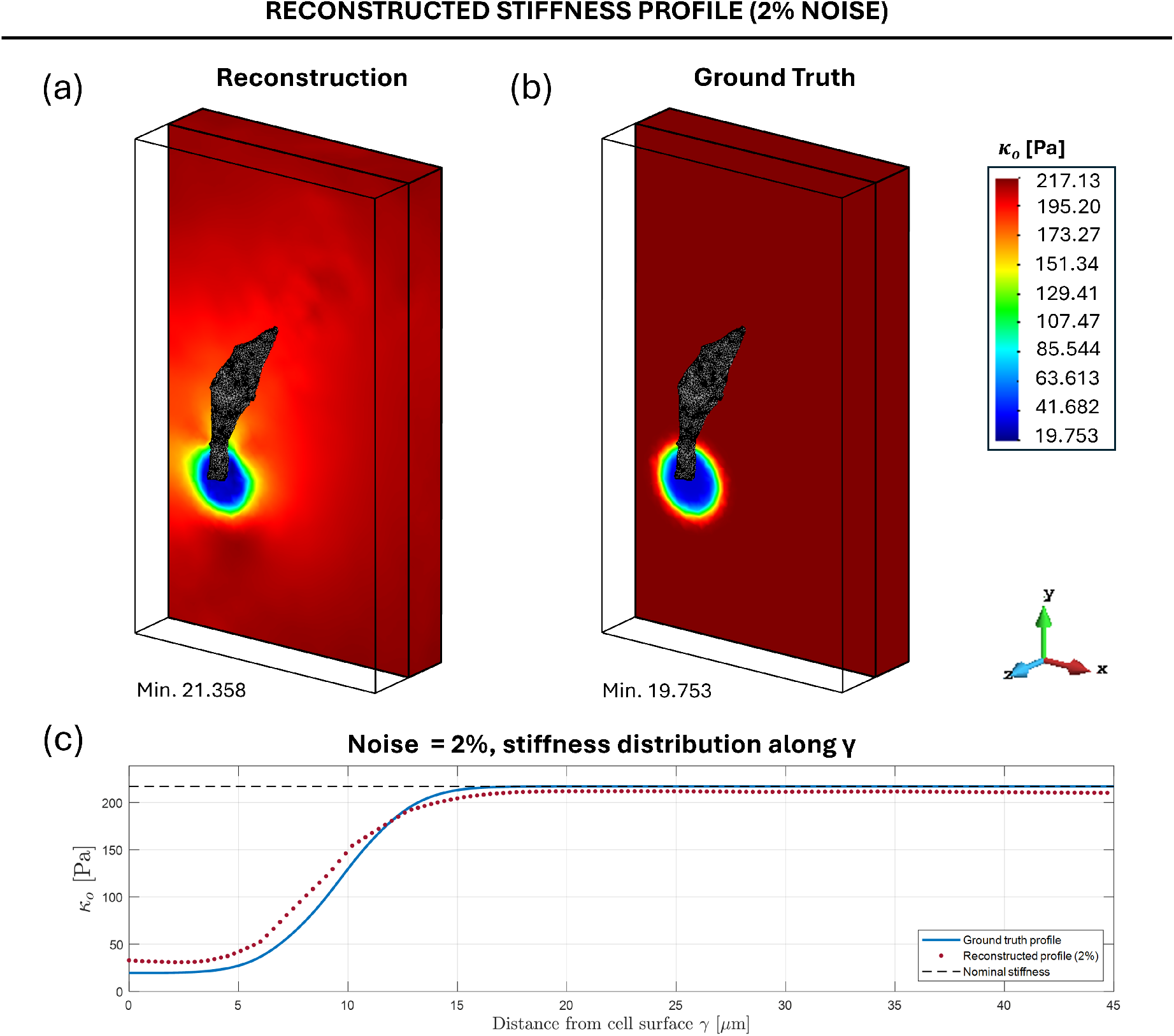
Material stiffness *κ*_*o*_ contours for the 2% noise case: (a) 3D view of mid cross-section of the hydrogel domain outlining reconstructed spatial material stiffness distribution. (b) 3D view of mid cross-section of the hydrogel domain outlining the ground truth spatial material stiffness distribution. (c) Reconstructed stiffness profile compared with the ground truth profile along *γ* (see Figure 3).

**Figure 8:**
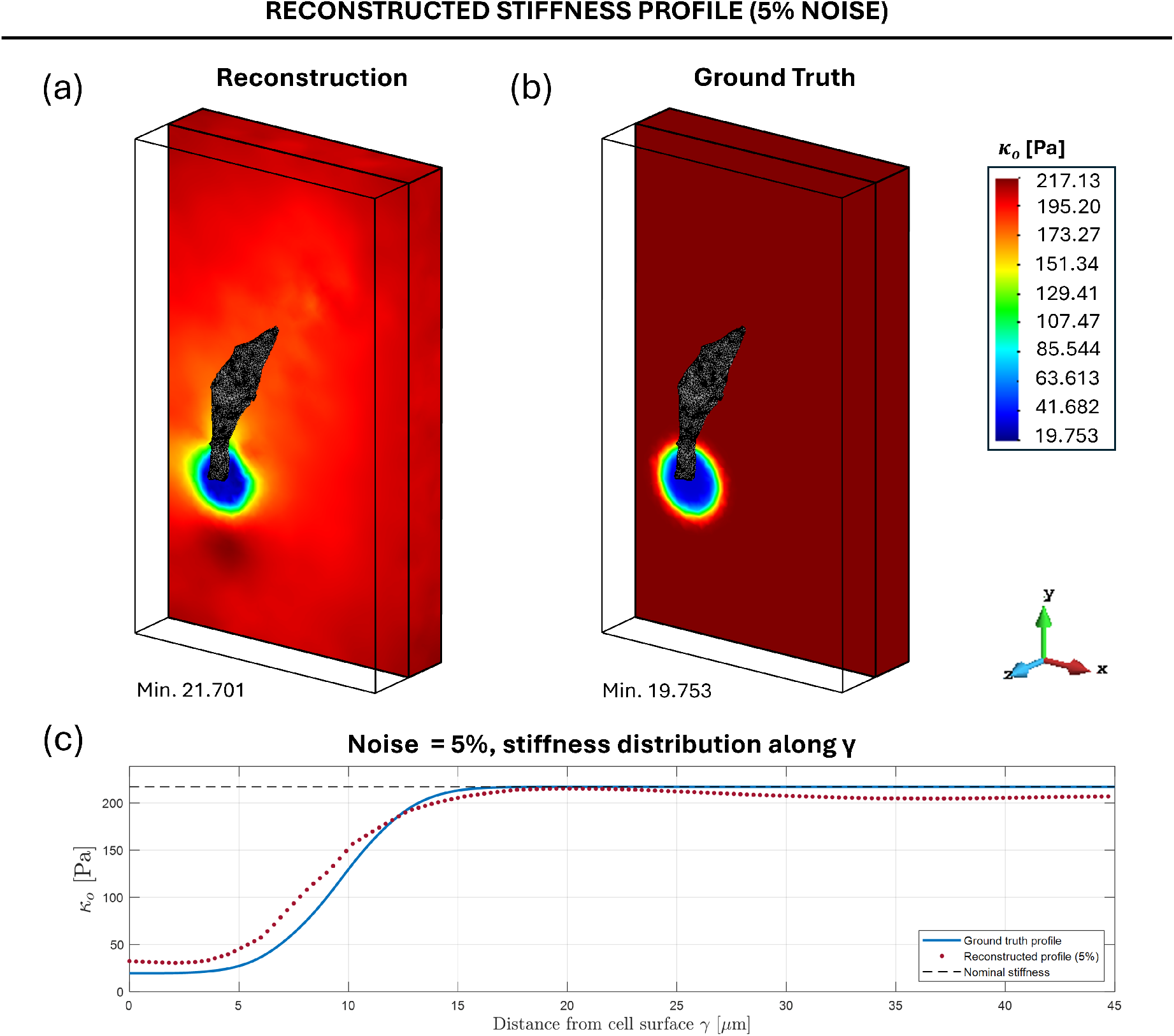
Material stiffness *κ*_*o*_ contours for the 5% noise case: (a) 3D view of mid cross-section of the hydrogel domain outlining reconstructed spatial material stiffness distribution. (b) 3D view of mid cross-section of the hydrogel domain outlining the ground truth spatial material stiffness distribution. (c) Reconstructed stiffness profile compared with the ground truth profile along *γ* (see Figure 3).

**Figure 9:**
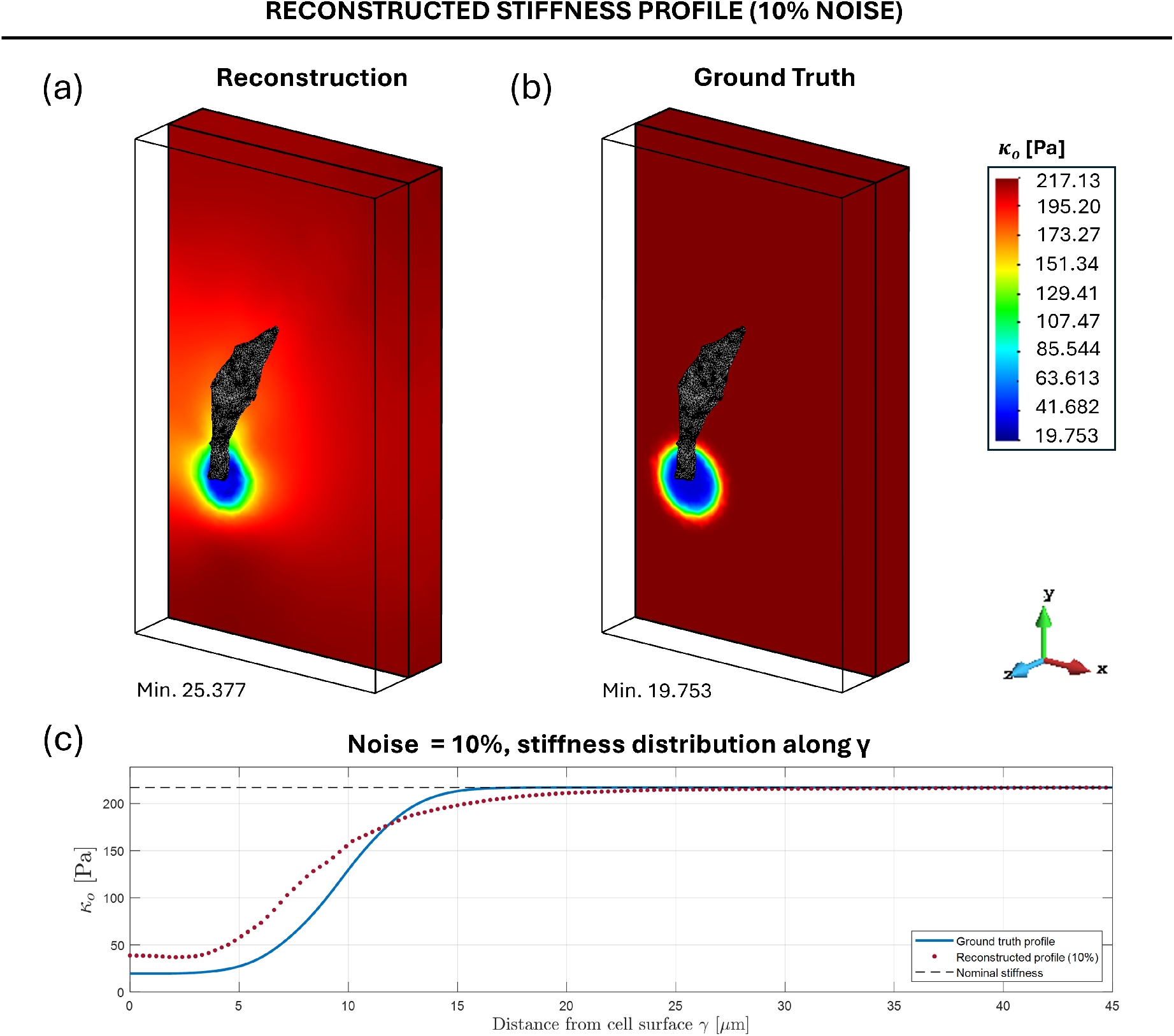
Material stiffness *κ*_*o*_ contours for the 10% noise case: (a) 3D view of mid cross-section of the hydrogel domain outlining reconstructed spatial material stiffness distribution. (b) 3D view of mid cross-section of the hydrogel domain outlining the ground truth spatial material stiffness distribution. (c) 3D stiffness profile compared with the ground truth profile along *γ* (see Figure 3).

The displacements magnitude contours presented in Figure 10 show excellent reconstructions for this variable, although the maximum displacement value is overestimated with respect to the ground truth case in all noise levels. Nonetheless, the low variation in displacement magnitude contours across the different noise levels show no clear tendency regarding overestimation of displacement values.

**Figure 10:**
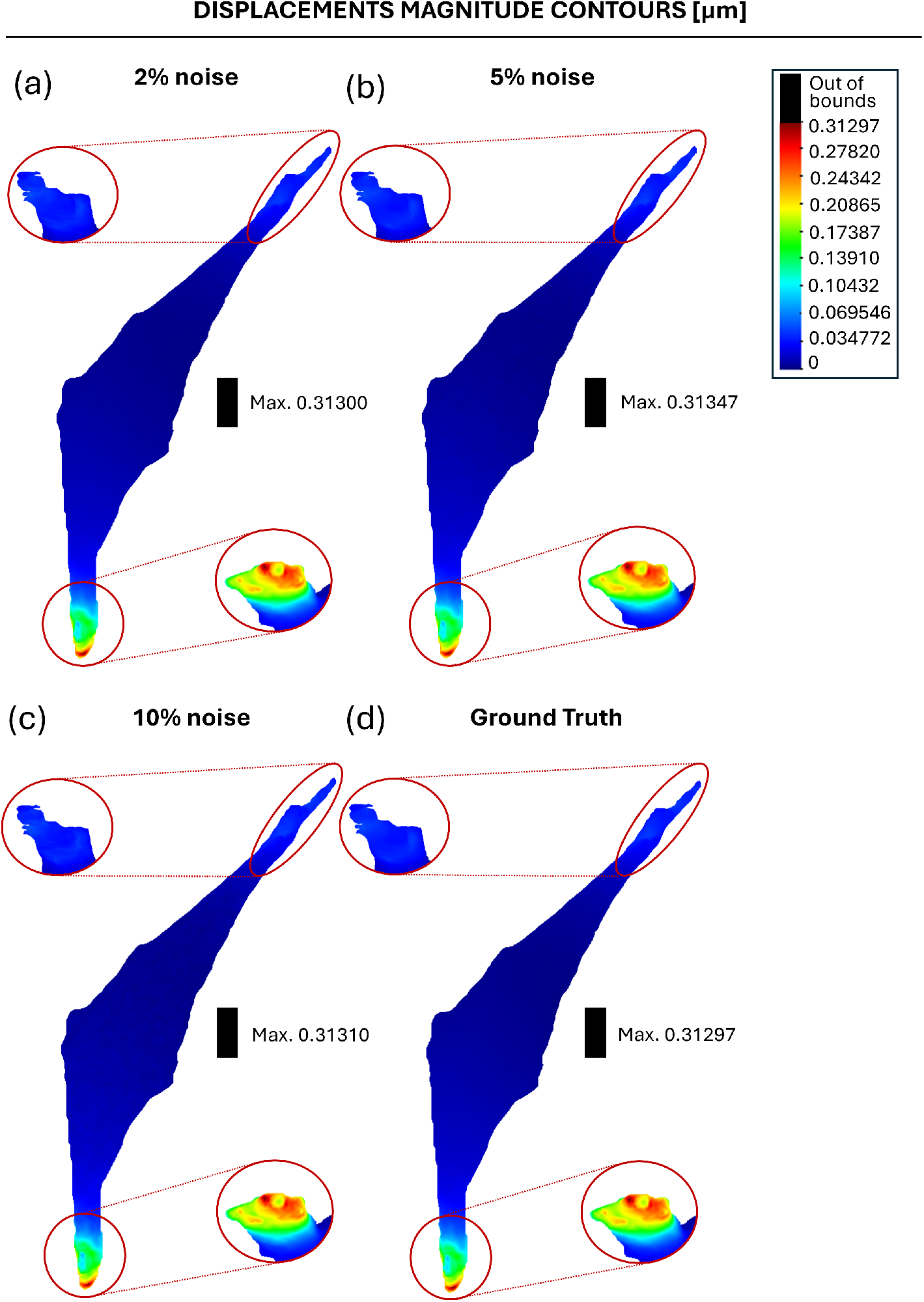
Displacements magnitude contours for each of the analyzed cases: (a) 2% noise. (b) 5% noise. (c) 10% noise. (d) Ground Truth solution.

Traction reconstruction is variably influenced by the degree of noise contained in the displacement data, depending on the magnitude of noise, as traction reconstruction is impacted by both the errors in reconstructing displacements and stiffness patterns. As it can be seen in Figure 11, traction profiles are well reconstructed for all noise levels, the general contours of the traction magnitude on the cell surface being overall very similar to those of the ground truth case (Figure 11d). However, the presence of noise in the measured displacements does influence the relative quality of these reconstructions, specially maximum tractions. Indeed, the maximum tractions are overestimated to some degree in all reconstruction cases, but this is emphasized in the third one (10%, Figure 11c). As a result, the traction profile becomes more difficult to accurately capture as noise increases, visually signalled by the cell surface regions colored in black where tractions have exceeded the lowest maximum value encountered within the cases, which corresponds to the ground truth case. Both the 2% and 5% noise cases (Figure 11a,b) show a fairly precise capture of the maximum traction value. Interestingly, for the latter, the maximum traction is less overestimated than that of the former. Nonetheless, in the 2% case, it can be appreciated how tractions acting on the main cell surface region (no protrusions) are much closer to the zero value of the ground truth case than in the 5% case (respectively 10%), rendering the reconstruction with 2% of noise the most accurate of the three. In light of this, displacement noise impacts traction reconstruction more specifically regarding both maximum and minimum tractions, as a result of accumulated errors in reconstruction of displacement and stiffness fields, whereas the overall distributions remain accurately reconstructed with respect to the ground truth case.

**Figure 11:**
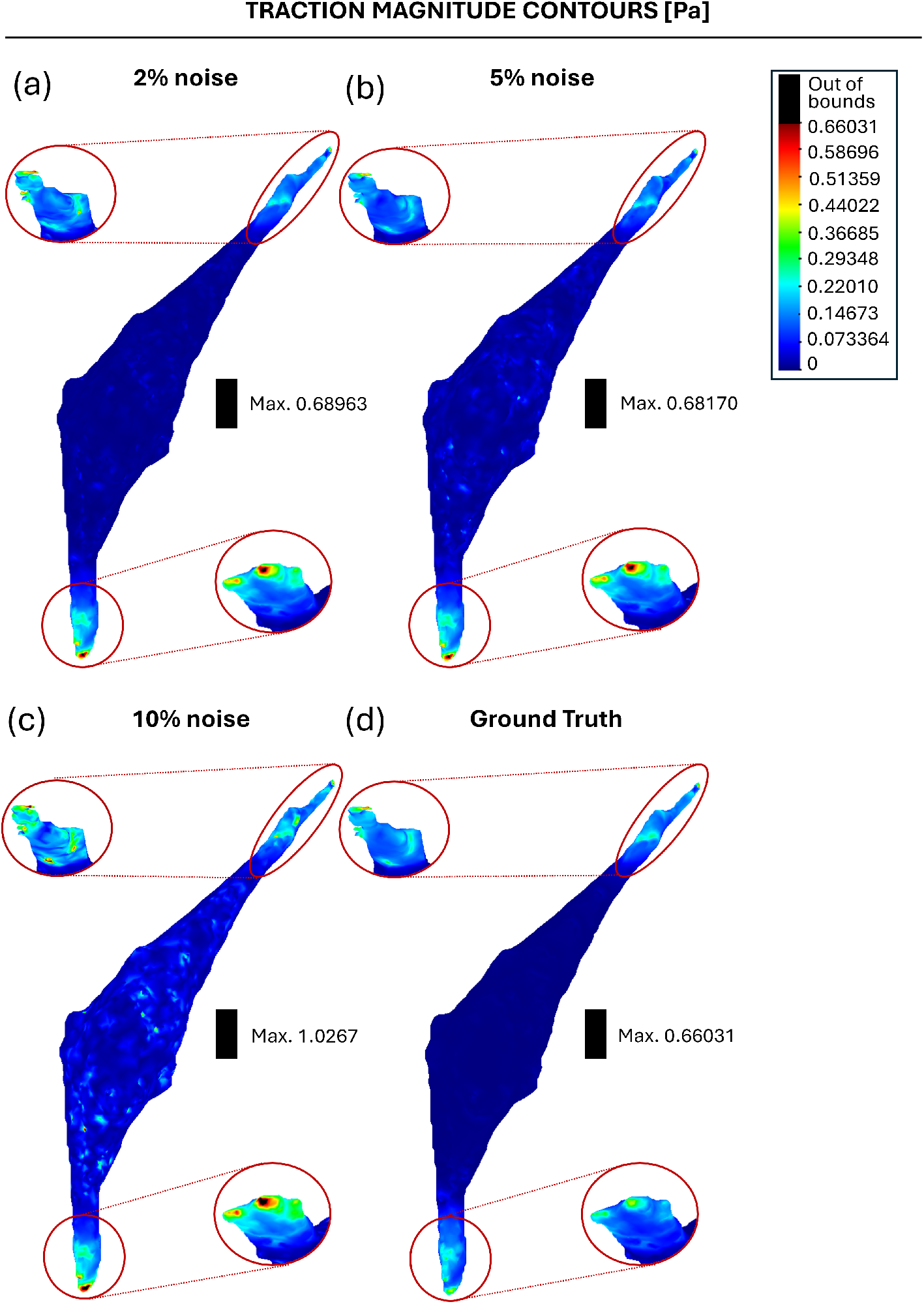
Traction magnitude contours for each of the analyzed cases: (a) 2% noise. 5% noise. (c) 10% noise. (d) Ground Truth solution.

## 5. Conclusions

In this paper, a novel inverse formulation devised to perform 3DTFM reconstructions with heterogeneous mechanical properties of the ECM is introduced. Using the (noisy) displacement field that is measured in a standard 3DTFM experiment, new displacements, tractions and the heterogeneous distribution of the hydrogel stiffness parameter can be retrieved. The mathematical details and the description of its computational implementation and general workflow are given. The methodology presents some novel aspects versus previous works. First, it is formulated within the context of nonlinear elasticity, and employs an adequate modelisation of the nonlinear behavior of collagen hydrogels. Also, the proposed methodology establishes a decomposition of a non-regularized constrained inverse problem into two coupled unconstrained regularized problems, which are combined in a NR/FEM computational framework. The reconstructions of the ground truth stiffness profile for two different degradation patterns, show that the NR/FEM implementation outperforms the L-BFGS iterative procedure, usually implemented in similar works, in regards to both accuracy and time of execution, particularly if a complete inverse reconstruction is required. Regarding the main recovered results from noisy displacement fields, it is shown that the method is able to provide good material stiffness reconstructions even for high levels of error added to the ground truth displacements solution. This is also the case for the reconstruction of tractions, which are overall well reconstructed in all cases. Nonetheless, the maximum tractions are found to be highly sensitive to the added noise magnitude when it exceeds a certain amount (the maximum concerned in this study). On the other hand, reconstructed displacement fields are qualitatively similar to the the ground truth case. It has ben also found that, added noise to displacements impacts the time of execution of the proposed methodology, increasing the number of local iterations taken until convergence. Specially, regarding the total number of local iterations taken by the inverse subproblem for the highest noise level case. Finally, the proposed methodology will be used to investigate *in vitro* 3DTFM experiments in which degradation of the ECM might be biologically relevant. As part of our future works, we intend to validate the results provided by the methodology with degradation measurements performed in the laboratory.

## Acknowledgements

A.A.-F. and J.A.S.-H. gratefully acknowledge the Financial support from MCIN/AEI/10.13039/501100011033 [PID2021-126051OB-C42]. The authors gratefully acknowledge Dr. Pablo Blázquez-Carmona and Raquel Ruiz-Mateos Brea for providing confocal image data of the cell geometry used in this study.

## Declarations

All authors declare that they have no conflicts of interest.

